# Polyploidy increases overall diversity despite higher turnover than diploids in the Brassicaceae

**DOI:** 10.1101/717306

**Authors:** Cristian Román-Palacios, Y. Franchesco Molina-Henao, Michael S. Barker

**Affiliations:** Department of Ecology and Evolutionary Biology, University of Arizona, Tucson, Arizona 85721, U.S.A.; Department of Organismic and Evolutionary Biology, Harvard University, Cambridge, Massachusetts 02138, U.S.A.; The Arnold Arboretum, Harvard University, Boston, Massachusetts 02131, U.S.A.; Departamento de Biología, Universidad del Valle, Cali, Valle 760032, Colombia

**Keywords:** Diversification, Evolution, Ploidy, Richness, Whole-genome duplications

## Abstract

Although polyploidy, or whole-genome duplication, is widespread across the Plant Tree of Life, its long-term evolutionary significance is still poorly understood. Here we examine the effects of polyploidy in driving macroevolutionary patterns within the angiosperm family Brassicaceae, a speciose clade exhibiting extensive inter-specific variation in chromosome numbers. We inferred ploidal levels from haploid chromosome numbers for 80% of species in the most comprehensive species-level chronogram for the Brassicaceae published to date. After evaluating a total of 54 phylogenetic models of diversification, we found that ploidy drives diversification rates across the Brassicaceae, with polyploids experiencing faster rates of speciation and extinction, but relatively slower rates of diversification. Nevertheless, diversification rates are, on average, positive for both polyploids and diploids. We also found that despite diversifying significantly slower than diploids, polyploids have played a significant role in driving present-day differences in species richness among clades. Overall, although most polyploids go extinct before sustainable populations are established, rare successful polyploids persist and significantly contribute to the long-term evolution of lineages. Our findings suggest that polyploidy has played a major role in shaping the long-term evolution of the Brassicaceae and highlight the importance of polyploidy in shaping present-day diversity patterns across the plant Tree of Life.

**Significance statement:** Although polyploidy is a source of innovation, its long-term evolutionary significance is still debated. Here we analyze the evolutionary role of polyploidy within the Brassicaceae, a diverse clade exhibiting extensive variation in chromosome numbers among species. We found that, although polyploids diversify slower than diploids, polyploids have faster extinction and speciation rates. Our results also suggest that polyploidy has played an important role in shaping present-day differences in species richness within the Brassicaceae, with potential implications in explaining diversity patterns across the plant Tree of Life.

## Introduction

Although polyploidy—the heritable condition of carrying more than two complete sets of chromosomes—is widespread across the plant phylogeny (1), its evolutionary significance is still debated (2-6). Discussions on the evolutionary role of polyploidy date back to Stebbins (7, 8) and Wagner (9) who considered polyploidy to have negligible effects on the long-term evolution of plants. This idea, later known as the ‘dead-end’ hypothesis, was recently reframed as an expectation that polyploid species will undergo extinction more frequently than diploids (4, 5, 10-12). However, polyploidy can influence the long-term evolution of lineages (1, 13-17) regardless of whether polyploids are more likely to go extinct than diploids (4,5, 10-12). Both interpretations of the ‘dead-end’ hypothesis are thus not equivalent to each other. The original hypothesis focused on the evolutionary role of polyploidy as a process (7-9), whereas the modern perspective compares macroevolutionary rates between polyploids and diploids (4, 5, 10-12). Discussions on the role of polyploidy in the evolution of plants have sometimes been obscured by the fact that there are different interpretations of the ‘dead-end’ hypothesis.

Here we evaluated the relationship between polyploidy, diversification rates, and clade richness in plants (7-9). Our analyses focused on the angiosperm family Brassicaceae, an ideal lineage for studying the short- and long-term evolutionary significance of polyploidy in flowering plants. The systematics and evolutionary history of Brassicaceae have been extensively studied (18-20). Furthermore, a comprehensive species-level phylogeny of the Brassicaceae including ∼48% of the extant species (1,667 out of ∼3,500 species) was recently published (21, 22). Haploid chromosome counts, which were used here to infer species ploidal levels, are available for ∼55% of extant species within the family (21-24). We used a model-based approach (25, 26) to infer ploidal levels for 1,336 species, representing more than 80% of the species in the tree and almost 40% of the total family richness. We then used the compiled ploidy database, along with the Brassicaceae phylogeny, to examine the role of ploidy in driving macroevolutionary rates within the family.

Using the Brassicaceae, we tested if polyploid species have significantly contributed to the short- and long-term evolution of plants, despite previous evidence that polyploids are more likely to go extinct than diploids (4, 5, 10-12). Although these previous studies compared the average net diversification rates of diploid and polyploid species, they did not examine the combined impacts of clade age and diversification at both ploidal levels on overall diversity and clade richness. Here, we used state of the art phylogenetic methods to test the relationship between diversification rates and ploidy. Specifically, we analyzed the fit of 24 different trait-dependent and trait-independent models of diversification that accounted for the potential effects of unassessed traits (i.e., hidden states; Hidden State Speciation and Extinction models; HiSSE; ref. 27) or not (Binary State Speciation and Extinction models; BiSSE; refs. 28, 29). We then examined the role of polyploidy in driving long-term evolution across clades by testing if polyploidy has significantly contributed to the present-day differences in species richness among clades. Using phylogenetic path regression analyses (30) of 65% of the total genera in the family (243 out of 372), we tested the contribution of polyploid richness on differences in species richness among clades in the Brassicaceae. We compared 30 models that assumed clade richness to be affected only by diversification rates and clade age to alternative models assuming ploidy influencing clade richness through net rates of diversification. With these species- and clade-level analyses in a large and well sampled phylogeny of Brassicaceae, we provide one of the most comprehensive tests of the relationship between polyploidy and plant diversity.

## Results

Our phylogenetic analyses of chromosome number evolution found that nearly half of extant Brassicaceae taxa have relatively recent polyploid origins. We collected haploid chromosome counts for 816 Brassicaceae species representing 49% of taxa sampled in the phylogeny (22) and 24% of the total family richness (Data S1; *Appendix SI*, Table S1; ref. 21). We used this database to infer ploidal levels for 1,333 species in the tree (80% of the species in the tree, and 38% of the total family richness; Fig. 1; *Appendix SI*, Table S2) using the likelihood-based approach implemented in ChromEvol (Data S2; refs. 24, 25). Overall, we inferred that nearly half of the analyzed Brassicaceae species were polyploids (*n=*654) and that these lineages were phylogenetically clustered in particular branches of the Brassicaceae phylogeny (Pagel’s lambda=0.816, P<0.001; D-statistic=0.006, P<0.001; Fig. 1). Alternative analyses of polyploid richness based on ploidy estimates from multiples of putative base numbers (“Stebbins fraction” sensu refs. 10,31) yielded largely similar results (see *Appendix SI*, Text S1).

**Fig. 1.**
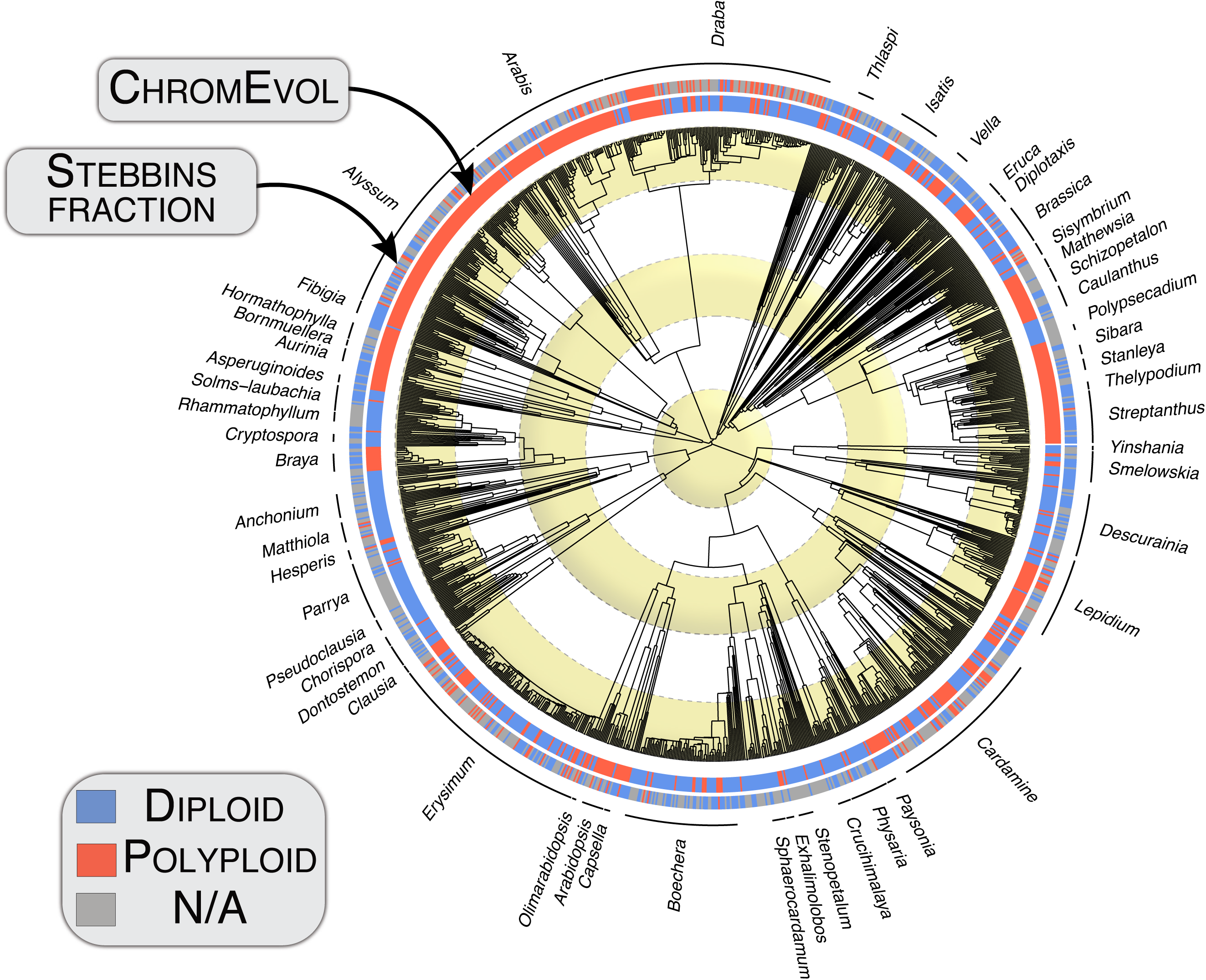
Phylogeny depicting the temporal evolution of ploidy across 1,366 Brassicaceae species (nearly 40% of the total family richness). Species are here coded as diploid or polyploid based on the evolution of haploid chromosome numbers across the tree. Ploidy levels were inferred using the Ploidy Inference Pipeline implemented in ChromEvol. We also show the ancestral state reconstruction of ploidy based on the best-fitting Hidden State Speciation and Extinction (HiSSE) model. Specifically, the favored HiSSE model assumed ploidy changes to influence diversification rates across the tree significantly. We depict also depict variation in net diversification rates across branches of the phylogeny – fast-evolving branches are marginally highlighted in black. Ancestral state reconstructions shown here are for visualization of diversification rates, and not used to ploidy changes across the tree. Instead, the ancestral state reconstruction for chromosomal numbers across the Brassicaceae is based on ChromEvol subtrees. Rings in the background are placed each ∼5 My.

Using ploidal inferences, we found that changes in ploidal level were important drivers of diversification rates across the Brassicaceae. We compared the fit of ten different trait-dependent diversification models that accounted for the potential effects of unassessed traits (*n=*6; hidden states; Hidden State Speciation and Extinction models, HiSSE; *Appendix SI*, Table S3; ref. 27) or not (*n=*4; Binary State Speciation and Extinction models, BiSSE; *Appendix SI*, Table S3; refs. 28, 29). The fit of each of these ten alternative models was then compared against two null models that assumed diversification rates to be independent of ploidy changes. Overall, based on the best-fitting HiSSE model (next model ΔAICc=9.786; *Appendix SI*, Table S3), we found that changes in ploidal level were important drivers of diversification rates across the Brassicaceae using either the ChromEvol or Stebbins Fraction estimates of species ploidal levels (Fig. 1; *Appendix SI*, Table S3).

Based on the best fitting HiSSE model (*Appendix SI*, Table S3), we estimate that polyploids, on average, speciate 48.7% faster (mean speciation rate polyploids=1.063 events My^−1^, diploids=0.715 events My^−1^; Fig. 2; *Appendix SI*, Table S4), and experience extinction 73.3% faster relative to diploids (mean extinction rate polyploids=0.863 events My^−1^, diploids=0.498 events My^−1^; Fig. 2; *Appendix SI*, Table S4). Congruently, we found that while diploids diversify 8.5% faster (mean diversification rate polyploids=0.199 events My^−1^, diploids=0.216 events My^−1^), evolutionary turnover is 58.8% faster in polyploids (mean turnover rate polyploids=1.926 events My^−1^, diploids=1.213 events My^−1^). In short, despite a slower mean rate of diversification for polyploid species compared to diploid species, the net rates of diversification in polyploids are, on average, positive.

**Fig. 2.**
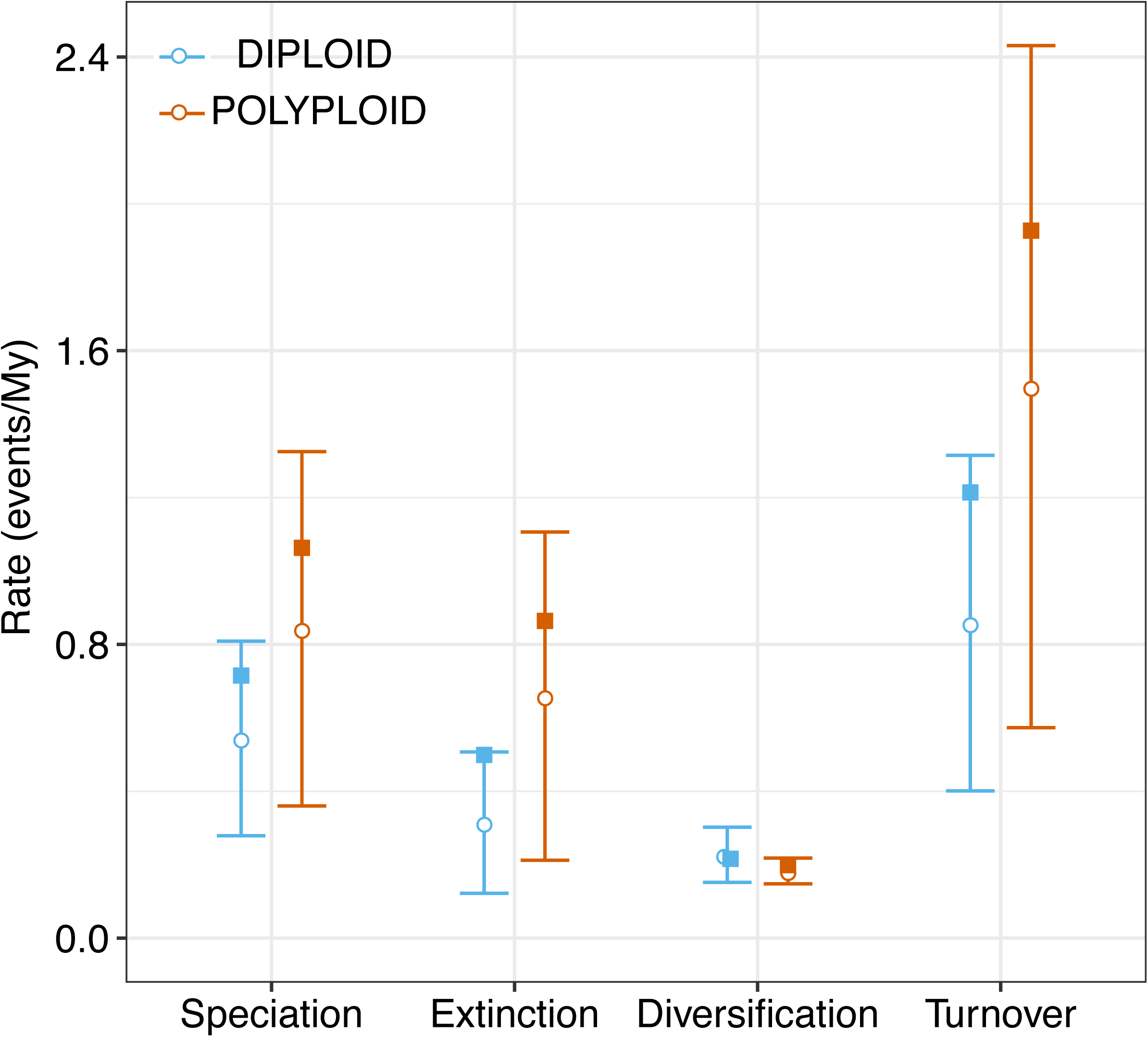
Macroevolutionary rates associated with polyploid and diploids states. We show speciation, extinction, diversification and turnover rates for each state based on HiSSE. Rates for observed states was estimated using the GetModelAveRates function in the hisse R package based on the marginal reconstruction of the best-fitting HiSSE model. We show median (circles), mean (squares), in addition to the 25% and 75% percentiles (lower and higher whiskers). Full results are presented in *Appendix SI*, Table S4.

Although these analyses provide insight into the different evolutionary dynamics of polyploid and diploid species, it is still unclear whether polyploidy-generated diversity significantly contributed to present-day differences in species richness across clades in the Brassicaceae. We used clade-level estimates of diversification rates to explore this question. These estimates of diversity were based on the Method-of-Moments estimator (MS hereafter; ref. 32) and the DR statistic (33). We note, however, that DR estimates have been shown to actually be closer to speciation and not diversification rates (34). Furthermore, the crown-based MS results excluded nearly 50% of genera in the phylogeny (see Methods). The main results we show, therefore, are for stem-based MS rates of diversification that include 97% of genera in the phylogeny and 65% of the total generic richness within the family (*Appendix SI*, Table S5).

To test whether polyploid richness influences species richness through diversification rates, we used phylogenetic generalized least squares models (PGLS; ref. 35). Strikingly, we found that 16–30% of the variance in net diversification rates was explained by polyploid richness alone (log-transformed polyploid richness versus net diversification rates; PGLS slope=0.085–0.444, r^2^=0.166–0.304, P<0.001; *Appendix SI*, Table S6). We also fitted PGLS regressions using alternative descriptors of polyploid richness within clades that were either unrelated to net diversification rates (log-transformed proportion of polyploid richness in described richness versus net diversification rates, PGLS r^2^=0.001–0.003, P>0.4; Table S6) or weakly associated (log-transformed proportion of polyploid richness in tree versus net diversification rates, r^2^=0.016–0.017, all P=0.04). Second, we used 14 phylogenetic path regression models (30,36) to test whether polyploid richness has indirectly shaped present-day patterns of species richness among clades through diversification rates (Fig. 3; *Appendix SI*, Table S7). Based on the best-fitting model—a model accounting for the indirect effects of diploid and polyploid richness and the direct effect of diversification and clade age on richness (next model ΔCICc=21.021; Fig. 3; *Appendix SI*, Table S7)—we found that polyploidy is an indirect driver of differences in species richness among clades across the Brassicaceae. Specifically, despite diploids contributing four times more than polyploids to net diversification rates, polyploidy-generated diversity has significantly influenced differences in species richness among clades through overall increased net diversification rates.

**Fig. 3.**
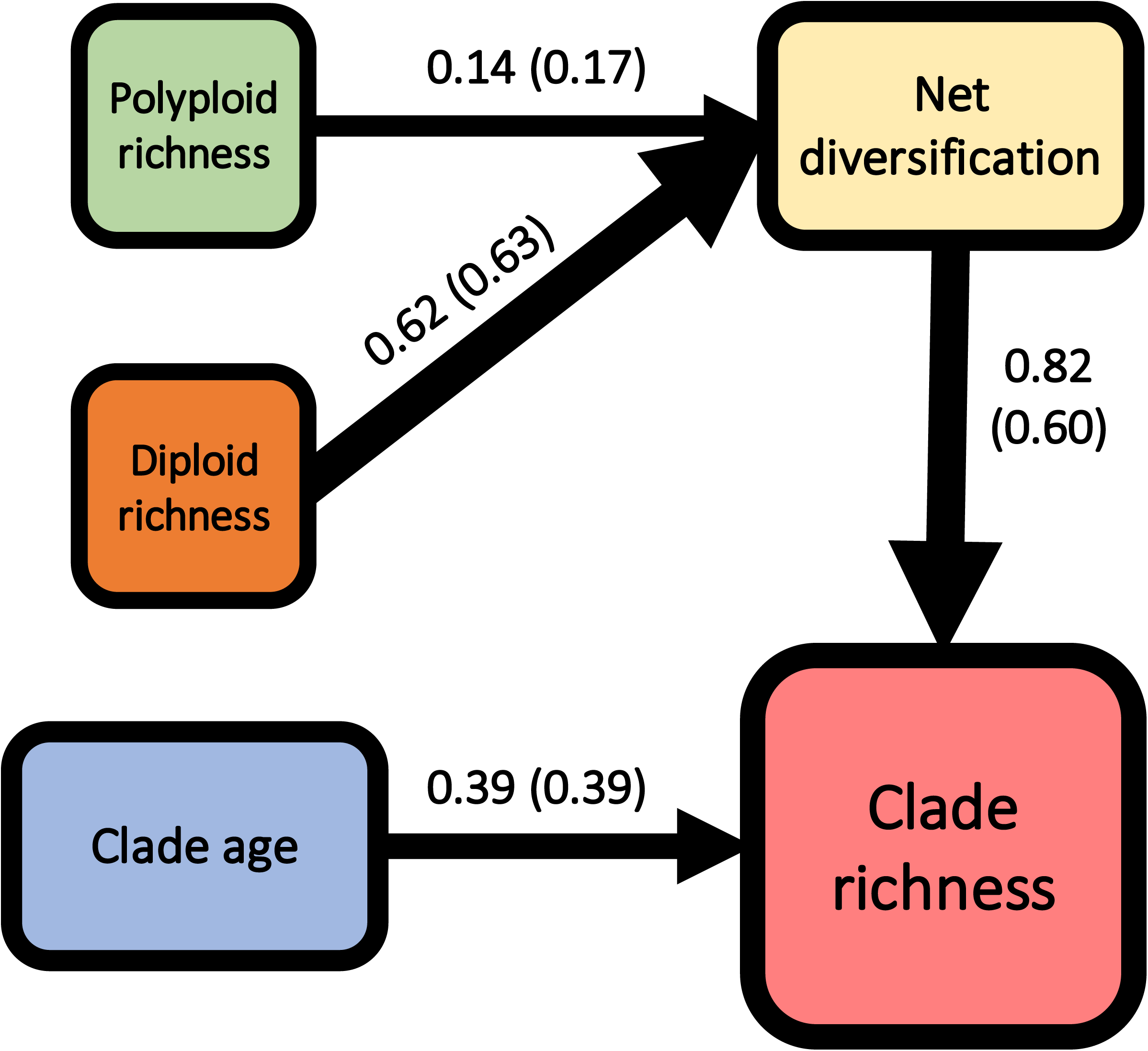
Phylogenetic path analysis showing the main drivers of differences in species richness among clades in the Brassicaceae. Results are based on 65% of the total generic richness within the family. We show results for the best-fitting phylogenetic path model. This model included the indirect effects of polyploid and diploid richness on species richness. We also note that the selected model assumed clade richness to be directly influenced by clade age and net rates of diversification. Here, polyploid richness is simply the number of polyploid species within each clade (based on ChromEvol analyses). We note that, contrary to the frequency of polyploidy, polyploid raw richness was a significant predictor of diversification rates explaining 16–30% of the variance. We present results for both stem-based MS and DR rates. We acknowledge that DR rates are usually referred to as related to speciation, but results are congruent to those based on MS. Path coefficient for the MS estimator is indicated below each arrow outside of parentheses. DR-related coefficients are indicated next to MS-related coefficients in parentheses. Finally, the clade age is always based on stem groups.

## Discussion

Our analyses revealed the complex interactions of polyploidy and diversification in flowering plants. Leveraging one of the largest species-level phylogenies for the Brassicaceae with ploidal level inferences for nearly 40% of all extant species from 65% of genera in the family, we found that polyploidy is not rare in the family. Consistent with previous estimates (10), we inferred that nearly half of extant Brassicaceae species are recent polyploids. The probabilistic inference of polyploidy using ChromEvol (24,25) estimated that 49% of species in the Brassicaceae were recent polyploids, whereas, the Stebbins fraction (10, 31) approach estimated that 26% of species were polyploid. These values are consistent with other estimates of polyploid incidence for angiosperms (4, 10, 37). A notable difference between these two estimates is that ChromEvol allowed us to infer ploidal level for species without chromosome counts. Regardless, the results of our phylogenetic analyses of polyploidy and diversification using either estimates were consistent with each other. Specifically, we found that polyploid species in the Brassicaceae experience higher turnover rates than diploids (4, 5). However, our deeper analyses of the differences in species richness among clades found that polyploidy significantly influenced present-day differences in diversity in the Brassicaceae. In particular, the best fitting model in a phylogenetic path analysis indicated that polyploid richness had a positive indirect effect on overall net diversification rates. Most previous research on this issue has focused only on the macroevolutionary differences between diploid and polyploid species (4, 10, 12, 38-40). Our analyses quantified the overall contributions of these differences to reveal that polyploidy has a direct, positive impact on overall net diversification rates and an indirect effect on differences in clade richness across the Brassicaceae.

If polyploidy has a positive effect on the diversity of the Brassicaceae, should it really be considered an evolutionary ‘dead-end’? The traditional perspective that polyploid plant species are evolutionary ‘dead-ends’ dates back to Stebbins (7, 8) and Wagner (9). These authors suggested that polyploidy in plants, although common, probably does not contribute significantly to the long-term evolution of lineages. However, recent studies (4, 5) have used this phrase to describe the macroevolutionary differences in extinction rates among polyploids and diploids. Although extinction rates are ultimately related to the long-term significance of polyploidy, the contribution of polyploid species to present-day diversity patterns should also be measured in terms of net diversification, turnover rates, and their actual effects on driving richness patterns across clades. Our finding that polyploidy positively influenced diversification rates and species richness within the Brassicaceae indicates that polyploidy is not an evolutionary ‘dead-end’ in the classical sense. Even in a family like the Brassicaceae where polyploid species have faster extinction rates than diploids, we still found that polyploidy has an overall positive effect on diversification and that polyploidy as a trait or process is not an evolutionary ‘dead-end.’

Despite our finding that the contribution of polyploid species to clade richness was significant, diploids contributed more than polyploids to variation in clade richness. This likely reflects the complexity of evolution and persistence at higher ploidal levels. Previous studies have discussed the evolutionary trade-offs associated with polyploidy (41-45). Specifically, polyploids tend to speciate faster than diploids but also to experience faster rates of extinction. On the one hand, faster speciation rates are potentially related to the rate of successful establishment of polyploids to novel habitats (46-55), or even to their ability to mask deleterious mutations (41-45, 56). On the other hand, polyploidy may trigger extinction rates by decreasing effective population sizes (57, 58), increasing rates of segregation errors during meiosis (59), or even by increasing the number of deleterious mutations that may contribute to extinction during diploidization (5, 58). The scale of our analyses may also play a role in the measured differences of net diversification rates between diploid and polyploid species. At deeper time scales above the family level, the number of polyploidy a lineage has experienced is positively correlated with increased diversification (6). Further, ancient polyploidy is correlated with mass extinction events and major geological transitions (59, 60), suggesting that polyploidy may facilitate lineage survival during periods of rapid ecological change. Our analyses within the Brassicaceae are not deep enough in time to capture that scale of ecological change. Further, the methods do not yet exist to formally estimate rates of polyploidy and diversification from inferences based on chromosome counts at the tips of a phylogeny with inferences of ancient WGD from genomic data deeper in a phylogeny. Progress on developing these types of integrated analytical frameworks and denser data sampling will permit further testing of the hypotheses on the role of WGDs in the evolution of plant diversity.

Our results recovering polyploidy as a major driver of diversification and species richness differences among clades shows that polyploidy is not an evolutionary ‘dead-end’ in the Brassicaceae. Our findings are potentially extended to other branches of the Plant Tree of Life that exhibit similar levels of polyploidy (e.g. Asteraceae, Poaceae, Solanaceae, Fabaceae, Cleomaceae; refs. 10, 13, 61-63) and ratios of polyploid and diploid net diversification rates (4). The influence of ploidy in the long-term evolution is potentially not only restricted to the Brassicaceae but likely to drive diversity in other lineages of green plants (6). Polyploidy has many impacts on the population genetics and ecology of species (5, 58, 64) that produce a complex association between WGD and macroevolutionary changes in diversity. Further quantitative studies are required to examine the specific relationship between diversification rates, clade richness, and ploidy across all green plants.

## Methods

### Brassicaceae phylogeny

The relationship between polyploidization and diversification rates is expected to be influenced, among others, by the phylogenetic relationships among species, and the evolutionary timing represented in branch lengths (27, 28, 65, 66). Because the phylogeny and timing of evolution in the Brassicaceae has been extensively discussed before (18-20, 67-71), we focused on analyzing the association between diversification rates and ploidy level changes across the phylogeny. Our analyses are based on the Brassicaceae subtree extracted using the extract.clade function (ape R package version 5.2; ref. 72) from the original GBMB Angiosperm phylogeny from Smith and Brown (22). The Brassicaceae subtree, including a total of 1,667 species (we pruned 100 tips corresponding to subspecies), is currently the largest-species level chronogram of the family. This phylogeny was constructed using PyPHLAWD based on GenBank release 218 (February 2017), with calibration points following Magallón et al. (73). Additional methodological details are summarized in Smith and Brown (22) and https://github.com/FePhyFoFum/big_seed_plant_trees. The Brassicaceae subtree analyzed here is provided in Data S1.

### Ploidy inference

Chromosome numbers for each species in the tree were retrieved from the Chromosome Count Database (CCDB; ref. 23) using the chromer R package version 0.1 (74). Raw haploid chromosome counts are provided in *Appendix SI*, Table S1. Species ploidal levels were then inferred with a maximum likelihood approach using the Ploidy Inference Pipeline implemented in ChromEvol v.2.0 (25, 26). Additionally, we also used a more conservative approach (10, 31) for estimating ploidal levels from haploid chromosome counts. Results for this alternative non-model based method were congruent with those based on ChromEvol and are presented in the supplemental results (see *Appendix SI*, Text S1).

Model-based ploidy inference analyses in ChromEvol failed to run on the full Brassicaceae phylogeny, consisting of 1,667 species, and with chromosome counts ranging between 4 and 156 (*Appendix SI*, Table S1). We note that the inference of species ploidy under ChromEvol represents a challenge given the size of the tree and variation in chromosomal counts (among and within) species. No study has used ChromEvol for analyzing chromosomal number evolution in phylogenies with more than 600 tips. Previous studies have analyzed trees containing between 18 and 588 species (4). To resolve this issue, we used an alternative approach to infer ploidal levels on our dataset (1,667 species). First, we estimated the median chromosome number across all the available counts for each species (see also ref. 75). This approach reduces the computational time required to run ChromEvol while summarizing the overall trend in chromosomal counts within each species. Second, we extracted all possible subtrees from the Brassicaceae phylogeny using the subtrees function implemented in the ape R package. We then selected subtrees with (i) sizes between 25 and 550 species, and (ii) a maximum of 70% of missing data (i.e., species lacking chromosome counts). This produced 342 subtrees that we ran the Ploidy Inference Pipeline (PIP) to infer ploidal level. The newly developed R function used to partition the Brassicaceae phylogeny into optimal subtrees for ChromEvol analyses is provided in Data S3. All ChromEvol-related files are provided in Data S2.

We used the default parameters for each Chromevol run (input files are provided in Data S2). We allowed ChromEvol to optimize the base number for each subtree. The PIP first compares the fit (AICc) among ten probabilistic models of chromosome number evolution along branches as a function of polyploidy and dysploidy (25). Confidence in ploidy assignment based on the best-fitting model is assessed from 100 simulations (76, 77). PIP classifies a taxon as polyploid if the maximum likelihood estimate of polyploidization from the root to the tip is larger than 0.9. Otherwise, PIP assigns a diploid state to the taxon. The PIP approach allowed us to infer ploidal levels for 517 species lacking chromosome counts. Unassigned species in our analyses were those with assignment probability lower than 0.9. Finally, we note that some authors refer to PIP-inferred polyploids as neopolyploids (77, 78). However, we refer to these lineages as polyploid given that the subtrees do not follow taxonomic limits (i.e., generic-level analyses as used in other studies).

Alternatively, we inferred species ploidy level relative to their generic base following Stebbins (31) and others (10, 79, 80)—an estimate sometimes called the “Stebbins’ fraction”. Specifically, we coded species as polyploid when the haploid count was greater than or equal to 3.5 times the lowest haploid (n) count of the corresponding genus (*Appendix SI*, Fig. S1, Table S2). However, the main results are based on ChromEvol given that (i) chromosome evolution in the Brassicaceae is faster than that assumed using the Stebbins’ fraction, (ii) Stebbins’ fraction depends on taxonomic delimitation, and (iii) ploidy cannot be estimated for species with missing data under the Stebbins method. For the latter reason, species-level sampling decreases to a half of analyzed richness under the ChromEvol database (i.e. 642 fewer species and 104 fewer genera than the database constructed using ChromEvol).

### Testing for phylogenetic signal of polyploidy

We analyzed the phylogenetic signal of ploidal level variation across the Brassicaceae phylogeny using Pagel’s Lambda (81) and the *D*-statistics (82). The first index measures the fit of the data to a Brownian motion model in which trait evolution matches the phylogeny. A lambda value close to one suggest a high phylogenetic signal in the trait, whereas values close to zero indicate that the trait is randomly distributed across the phylogeny (81). We used the phylosig function in Phytools version 0.6-60 (83) to estimate the Lambda value of ploidal variation across the tree. Statistical significance was approached on 1,000 simulations. We also used the *D*-statistics to measure the phylogenetic signal of binary traits (such as polyploid vs diploid). We estimated the phylogenetic signal under the *D*-statistics using the phylo.d function in the Caper R package version 1.0.1 (84). Values of *D*-statistic close to 0 indicate the trait conservatism expected under Brownian motion, and a value of 1 indicates a random distribution of the trait across the tree (82). For both analyses, the ploidy database is provided in *Appendix SI*, Table S2 and the phylogeny in Data S4.

### Species-level diversification analyses

We used both Binary State Speciation and Extinction models (BiSSE; 28, 29) and Hidden State Speciation and Extinction models (HiSSE; 27) to examine the importance of ploidy changes on diversification of the Brassicaceae. These models also allowed us to quantify rates of speciation and extinction associated with polyploid and diploid states. BiSSE models were fitted using the diversitree R package version 0.9-10 (85). HiSSE models were fitted using the hisse R package version 1.8.9 (27, 66). Species were coded as being either diploid (0) or polyploid (1) based both on the Stebbins Fraction and ChromEvol-inferred ploidy states. We specified the following sampling fraction in both BiSSE and HiSSE models based on the ploidy database generated using ChromEvol: diploid=0.499, polyploid=0.501 (*Appendix SI*, Table S2).

We fitted a total of 12 different trait-dependent diversification models (both BiSSE and HiSSE; *Appendix SI*, Table S3). Four of the analyzed models corresponded to a hypothesis of trait-dependent diversification under a BiSSE framework. Using the hisse function in the hisse R package, we first fitted a model assuming turnover, extinction, and transition rates to be constrained between states (i.e., diploid and polyploid). A second model constrained only transition rates between states, with turnover and extinctions free between states. A third model constrained both turnover and extinction rates to be equal between states but allowed transition rates to vary between states. Finally, we fitted a model allowing all parameters to be free between states (turnover, extinction, and transition rates).

Next, we focused on fitting eight additional models under a HiSSE framework (*Appendix SI*, Table S3). We first fitted two-character independent models (CID2, CID4) that assumed diversification rates to be exclusively driven by the hidden states (states A or B; not by the observed states 0 or 1). We fitted a 2-state character independent model using the hisse function in the hisse package. This null model has the same number of parameters as a BiSSE model (see above; ref. 27). We specified only two free parameters in the turnover.anc and eps.anc arguments of the hisse function (one parameter for 0A and 1A, and another parameter for 0B and 1B). Alternatively, we fitted the 4-state character independent model under default parameters using the hisse.null4 in the hisse R package. This null model has the same complexity as a general HiSSE model. The remaining six HiSSE models correspond to alternative hypotheses of trait-dependent diversification. These models were fitted using the hisse function implemented in the hisse R package. Specifically, we first fitted a model with two hidden states being present, independent turnover and extinction parameters across the four states, and transition rates constrained to be equal. The second and third HiSSE models assumed only a single hidden state (hidden state present in either state 0 or 1), with turnover and extinction rates set to be free among states. Transition rates were constrained to be equal between states. HiSSE models four and five assumed only a single hidden state for each observed state (hidden state for either state 0 or 1). All the remaining parameters (turnover, epsilon, and transition rates) were set free between states. Finally, we fitted a full HiSSE model. This model included two hidden states, assumed independent diversification parameters across states, and allowed for different transition rates among states.

We compared the fit among the analyzed 12 models using Akaike Information Criteria (AIC values: dAIC, and wAIC; *Appendix SI*, Table S3; ref. 86). Akaike weights were estimated using the akaike.weights in the qpcR R package version 1.4-1 (87). We then focused on analyzing the outcome of the best fitting model(s). For this, we used the MarginRecon (hisse R package) to visualize the association between ploidy changes and diversification rates across the Brassicaceae phylogeny. We also estimated the confidence intervals for speciation and extinction rates associated with each state in the best fitting model(s) using the SupportRegion implemented in the hisse R package. Finally, we summarized rates of speciation, extinction, and net diversification at the tips of the tree by running the GetModelAveRates (hisse R package) on the marginal reconstruction object from the best-fitting model. The species-level phylogeny analyzed in HiSSE and BiSSE is provided in Data S4. Similarly, the ploidy dataset is provided in *Appendix SI*, Table S2.

Finally, the same set of analyses were repeated for ploidal levels inferred using the Stebbins fraction. Results, which were congruent to the ones based on ChromEvol, are presented in *Appendix SI*, Table S3, and *Appendix SI*, Text S1.

### Clade-level diversification analyses

We analyzed the role of polyploid species in driving species richness among Brassicaceae genera using phylogenetic regressions (35) and phylogenetic path analyses (30). We first constructed an generic-level phylogeny for the family by pruning from each genus in tree, all tips in each genus except one. This step was conducted using the drop.tip function in the ape R package.

We first constructed a database summarizing clade-level diversification rates (two different estimators; see below), species richness, clade age (stem and crown), and the proportion of polyploid species within clades. We retrieved species richness for each of the 243 analyzed genera from (21). We estimated rates of diversification within each genus using the Method-of-Moments estimator (MS hereafter; 32). Net diversification was estimated for the crown and stem groups after assuming three different relative extinction fractions (e=0, e=0.5, and e=0.9). MS rates were estimated using the bd.ms function implemented in the geiger R package version 2.0.6.1 (88,89). We note that 97% of genera in the tree were monophyletic (243 out of the total 250). The remaining 3% non-monophyletic genera (n=7) were simply excluded from the phylogenetic path analyses. We also highlight that 51% of Brassicaceae genera were excluded from crown-based analyses. These lineages were represented by a single species in the tree (124 of 243). Stem-based analyses are, however, based on all the 243 analyzed genera.

Alternatively, we estimated species-specific rates of diversification (potentially reflecting speciation rates; refs. 34, 90, 91) based on the DR statistic (33). We estimated species-specific rates based on the full species-level phylogeny of the Brassicaceae (1,667 species). Species-specific DR rates, summarized in *Appendix SI*, Table S9, were then used to estimate the mean DR within genera (*n=*243).

We summarized the richness of polyploids within each clade using three different indexes. First, we simply obtained the absolute number of polyploid species within each genus (log-transformed). Second, we estimated the ratio between polyploid richness within each clade and the total number of species sampled in the tree for the same genus. Third, we estimated the ratio between polyploid richness and total clade richness. The resulting generic-level tree is provided in Data S5. The analyzed database including clade age, richness, diversification rates, and proportion of polyploids is provided in *Appendix SI*, Table S5. Then, we tested whether diversification rates among genera was influenced by polyploidy. We specifically tested the relationship between MS and DR rates against each of the three indexes of polyploid richness (see above). Phylogenetic regressions (35) were fitted using the caper R package. We used the pgls function from the same package and allowed lambda to be estimated from the dataset.

Finally, phylogenetic path analyses were used to test the indirect effect of polyploidy in species richness among clades. The following procedure was repeated for each polyploidy index that was found to significantly predict diversification rates (see above PGLS). Models were fitted using the define_model_set function implemented in the R phylopath package version 1.0.2 (36). Models were analyzed in phylopath package under a lambda model of evolution developed to analyze continuous traits. We fitted and compared a total of 30 phylogenetic path models (25 MS-based models, and five DR-based) that tested the indirect contribution of polyploidy in species richness among Brassicaceae genera. However, the main results are presented for stem-based rates of net diversification based on MS estimator (see below). Results were largely congruent to those based on DR rates because both approaches included all the 243 analyzed genera. Phylogenetic path regression models assumed different clade age (stem, crown), diversification rate estimates (MS based on crowns or stems, or DR), and variable influence of polyploid and diploid richness on net diversification rates. Our main results are, therefore, based on both stem-based MS and DR (all clades are included). Models were compared using a modified version of the Akaike Information Criteria (AIC; ref. 86) that was developed for phylogenetic path analyses. This index is known as the C statistic Information Criterion (i.e., CIC statistic; ref. 30) and is also calculated in the phylopath R package.

We note that some of the fitted phylogenetic path models tested the association between clade richness and polyploid richness. The positive association between polyploid richness and species richness has been demonstrated using both theoretical and empirical data (58, 92-93). Nevertheless, we note that our results are not affected by the “inertia” in polyploid richness. First, previous analyses on the relationship between species richness and polyploid richness assumed clades to be independent of each other. However, the fact that lineages share an evolutionary history was not accounted in previous studies. Second, in addition to the shared evolutionary history, we accounted for the effect of clade age in the association between polyploid richness and clade richness. The potential temporal scaling in the association between clade richness and polyploid diversity was not accounted for by previous studies (94, 95).

The association between clade richness and polyploid richness may also be driven by the differential contribution of allo- and auto-polyploids to the overall polyploid richness. Specifically, one would expect allopolyploid richness to be positively associated with clade richness given that allopolyploids lineages are potentially more likely to be formed in species-rich clades. However, it is also known that successful allopolyploids are more likely to be formed by distantly related diploids than closely related ones (96, 97). Autopolyploidy, on the other hand, is potentially unrelated to clade richness. Our analyses based on chromosomal counts cannot differentiate between allo- and autopolyploids. However, models that assumed polyploid richness to be dependent on clade richness fitted worse than other models. It is thus unlikely that the association between polyploid richness and clade richness to be an artifact of the predicted diversity-dependence richness in allopolyploids.

Finally, we highlight that missing data have a significant effect on the inferred effects of ploidy on species richness (through net diversification rates). Specifically, our results were congruent when DR and stem-based MS rates were analyzed for 243 of the genera, but crown-based results for MS estimator, where 119 clades were excluded, were remarkably different. Given the effect of missing data on the resulting patterns, we did not perform phylogenetic path analyses based on the Stebbins Fraction dataset. Phylogenetic path analyses cannot account for missing data. Specifically, the Stebbins Fraction ploidal level dataset included only 138 genera, which represents 55% of the genera in the tree, and 37% of the total family diversity.

## Data availability

All data and code are available in the manuscript, supplementary materials. Data S1–S5 are available in FigShare (https://figshare.com/s/cea330feaca5bc584b32).

## Acknowledgments

We thank Jeremy Beaulieu for help with HiSSE, Zheng Li, and Elizabeth Miller for discussions, and Elena Kramer for providing comments on a previous versions of the manuscript. Hosting infrastructure and services were provided by the Biotechnology Computing Facility at the University of Arizona and FAS Research Computing Facility at Harvard University. Y.F.M.H. was supported by a Graduate Fellowship from the Department of Organismic and Evolutionary Biology at Harvard University; the Colombian Administrative Department of Science, Technology, and Innovation (COLCIENCIAS) 538-2012; and the Fulbright Exchange Visitor Program G-1-12218. M.S.B. was supported by National Science Foundation (NSF) IOS-1339156 and NSF EF-1550838.

## Author contributions

C.R.P., Y.F.M.H., and M.S.B. designed research; C.R.P., and Y.F.M.H., performed research and analyzed data; and C.R.P., Y.F.M.H., and M.S.B wrote the paper.

## Notes

https://figshare.com/s/cea330feaca5bc584b32

